# Labeling RNAs in live cells using malachite green aptamer scaffolds as fluorescent probes

**DOI:** 10.1101/173898

**Authors:** V. Siddartha Yerramilli, Kyung Hyuk Kim

**Affiliations:** Department of Bioengineering, University of Washington, Seattle, Washington 98105, United States

**Keywords:** RNA aptamer, malachite green, scaffold, synthetic biology, live cell imaging

## Abstract

RNAs mediate many different processes that are central to cellular function. The ability to quantify or image RNAs in live cells is very useful in elucidating such functions of RNA. RNA aptamer-fluorogen systems have been increasingly used in labeling RNAs in live cells. Here, we use the malachite green aptamer (MGA), an RNA aptamer that can specifically bind to malachite green (MG) dye and induces it to emit far-red fluorescence signals. Previous studies on MGA showed a potential for the use of MGA for genetically tagging other RNA molecules in live cells. However, these studies also exhibited low fluorescence signals and high background noise. Here we constructed and tested RNA scaffolds containing multiple tandem repeats of MGA as a strategy to increase the brightness of the MGA aptamer-fluorogen system as well as to make the system fluoresce when tagging various RNA molecules, in live cells. We demonstrate that our MGA scaffolds can induce fluorescence signals by up to ~20 fold compared to the basal level as a genetic tag for other RNA molecules. We also show that our scaffolds function reliably as genetically-encoded fluorescent tags for mRNAs of fluorescent proteins and other RNA aptamers.

---

RNAs mediate many essential processes in live cells, both as the messenger RNA (mRNA) that transcribes genetic information to protein and as non-coding RNAs that regulate gene expression at the transcriptional and post-transcriptional level(1,2). A high degree of non-homogeneous RNA localization occurs typically within different intracellular compartments(3,4). This heterogeneous distribution of RNA contributes greatly to the complexity in studying RNA in live cells. Conventional fluorescence techniques such as fluorescence in situ hybridization (FISH) that involve damaging the natural structure of the cell due to fixation provide high resolution images of RNA molecules(5), but are not appropriate to measure the dynamics of RNA in live cells. Hence fluorescent tagging of RNA can provide a major advantage of observing the spatio-temporal distribution of RNA over standard biochemical methods. The MS2 system is one such example that uses the strong and specific binding of a bacteriophage protein MS2 to a short RNA sequence(6-9). Fluorescent proteins such as GFP are tagged to MS2 and the RNA recognition motif is placed at the 3’ end of an mRNA sequence. However, this approach requires pre-expression of high levels of GFP resulting in strong background signals that makes the measurement of RNA concentrations at the cellular level difficult unless specific measures are taken to fix or localize it(10).

Fluorescence-activating or light-up aptamers are small genetically encoded RNA molecules whose binding to their ligands results in the production of fluorescent signals(11,12). RNA aptamers can be applied to potentially quantify RNA expression and transcription dynamics within live cells. Malachite green aptamer (MGA) is one of the first fluorescence-activating aptamers to be described in the literature(12,13). It was identified through *in-vitro* selection for its high binding affinity and strong fluorescence intensity. It binds to malachite green (MG), a commercially available triphenylmethane dye that strongly absorbs light energy at 618 nm (12) dissipating it by heat through rotational movement of its phenyl groups. When bound to MGA, the restriction of MG molecules’ movement shifts their absorbance from 618 to 630 nm, resulting in the emission of far-red fluorescence at 655 nm (12,13).

MG can easily penetrate bacterial cell membranes, while eukaryotic membrane permeability is enhanced by ester linkages(14). Because of its properties, MGA has been one of the aptamer candidate systems that has the potential to be applied in *in-vivo* applications. However, initial studies have shown that aptamer fluorescence in bacteria was only barely above cellular autofluorescence level, possibly due to high degradation and improper folding(15). One strategy to enhance the fluorescence signals was to stabilize MGA by encasing it within a 5S RNA (MGA-5S). One study showed preliminary findings where MGA fluorescence was seen *in-vitro* and in *E. coli* using MGA-5S(16). MGA-5S fluorescence in *E. coli* was further explored in another study where an inducible MGA-5S model showed up to a 3-fold increase in batch culture and up to a 10-fold when cultured in a chemostat(15). This study also showed a concurrent expression of MGA and GFP fluorescence using a fusion RNA only when certain linkers were used to separate MGA-5S and the GFP mRNA(15).

Studies (8,9) have shown that adding many RNA repeats significantly enhanced fluorescence in bacterial and yeast cells using the MS2 systems. This approach of using RNA repeats has also been applied to enhance aptamer fluorescence using Spinach aptamer(17). It was seen that expression of a scaffold or array of multiple Spinach aptamer copies in bacterial cells enhanced Spinach-DFHBI (3,5-difluoro-4-hydroxybenzylidene imidazolinone) fluorescence as detected by imaging. Increasing the number of Spinach repeats resulted in a non-linear increase in aptamer-induced fluorescence intensity in cells upon being expressed. The expression of 16 Spinach repeats showing a 4.5 fold increase in fluorescence intensity compared to single Spinach RNA, while 64 Spinach repeats showing a 11-fold increase(17). Therefore, expressing a scaffold of multiple MGA repeats in tandem can, in principle, solve the issue of low fluorescence and high background *invivo*. A recent study using malachite green aptamer scaffold also showed increased fluorescence *in-vit*ro but not *in-vivo* in mammalian and yeast cells(18).

Here, we explore the expression of a scaffold consisting of 6 copies of MGA in bacterial cells, showing a compatible spectrum of fluorescence with other currently available RNA aptamers. We observe a strong and stable MGA fluorescence that can be linked to multiple kinds of RNA molecules without significant fluorescent signal loss in *E. coli*. This MGA fluorescence can be studied using microscopy and flow cytometry showing its potential as a tool to study RNA expression and dynamics *in-vivo*.

## Results

### MGA fluorescence is enhanced by aptamer scaffolds

We designed scaffolds of MGAs with various repeats that are separated by small linkers. The MGA scaffolds were integrated into a pET28a plasmid, which allowed MGA’s *in-vitro* and cellular expression to be regulated by the T7 promoter and lac operator system respectively. First, six repeats of MGAs (MGA6) were designed by joining six MGAs with five distinct linkers, and then twelve repeats of MGAs (MGA12) were constructed by joining two different MGA6 scaffolds. MGA4 was produced by deletion of two different MGA structures. MGA-5S – a single MGA aptamer enclosed in a 5S ribosome subunit that has been used in prior work(15,16) – was also constructed in a pET28a plasmid. The plasmids were then transfected into BL21DE3 strain of *E. coli*.

For all studies, MGA expression was induced by 0.1 mM Isopropyl β-D-1-thiogalactopyranoside (IPTG) and quantified immediately after the addition of 5µM MG. *E. coli* containing MGA6-pET28a and MGA12-pET28a, respectively, showed a very strong MGA fluorescent signal upon induction more than 20-fold higher than that of control (Figure 1A). Cells containing MGA4-pET28a and MGA-5S-pET28a also showed a moderately strong MGA fluorescence upon induction, which was slightly smaller than that of MGA6 or MGA12. The distribution function of the MGA fluorescence signals at the single cell level was obtained (Figure 1B) and the cell-to-cell variability in the fluorescence signals was quite strong (coefficient of variation for the induced sample =1.0). The induced MGA fluorescence was also seen by imaging using a microscope in the presence of MG (Figure 1C).

**Figure 1:**
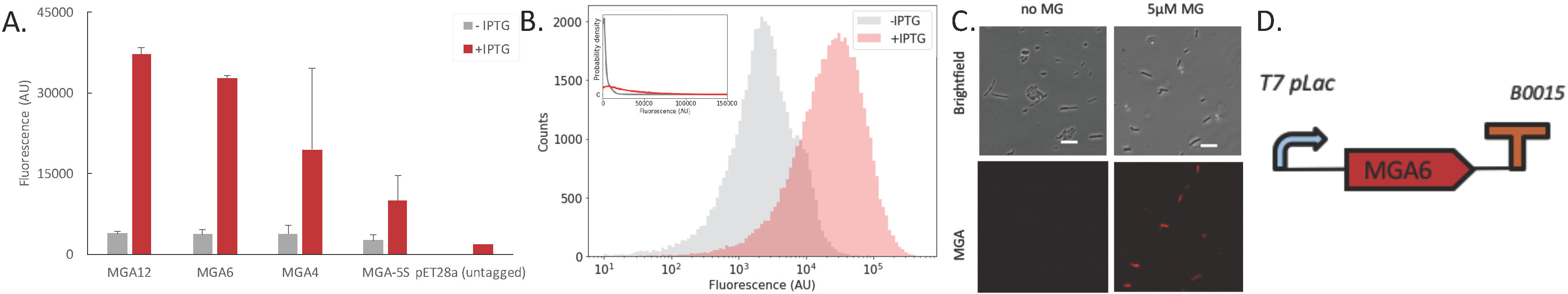
Inducible malachite green aptamer expression. (A) MGA fluorescence in induced (+IPTG) and uninduced (- IPTG) *E. coli* BL21DE3 cells containing a pet28a plasmid consisting of a MGA12, MGA6, MGA4 or MGA-5S scaffold was measured in a flow cytometer. pET28a (untagged) is a control that does not have any aptamer sequence in the ORF. (B and C) MGA fluorescence in individual cells containing MGA6-pet28a was measured via flow cytometry and observed using a microscope. MGA expression was induced by adding 0.1 mM IPTG to cells growing in M9CA media (Materials and Methods). MGA fluorescence is seen in response to 5 µM of malachite green. The scalebar in (C) represents 10 µm. SBOL compliant figure of the construct is shown (31).

The fluorescence signals of the MGA scaffolds can be enhanced further by optimizing cell density in the culture and by treating the culture with rifampicin. Our flow cytometry analysis has shown that the strength of the fluorescence signals per cell is positively correlated with cell density in the culture (Figure S1). This may be related to the fact that the cells grow more slowly in dense culture and leads to slower dilution of the MGA caused by slower cell doubling, while MGA synthesis is kept roughly the same. Another way to enhance the signals is by rifampicin treatment. Rifampicin, an inhibitor of bacterial DNA-dependent RNA polymerase, does not inhibit the activity of T7 polymerase (22). Thus, it can make the transcription cellular machinery more dedicated to T7 polymerase activity. Figure 2 shows that MGA fluorescence was increased significantly after rifampicin treatment.

**Figure 2:**
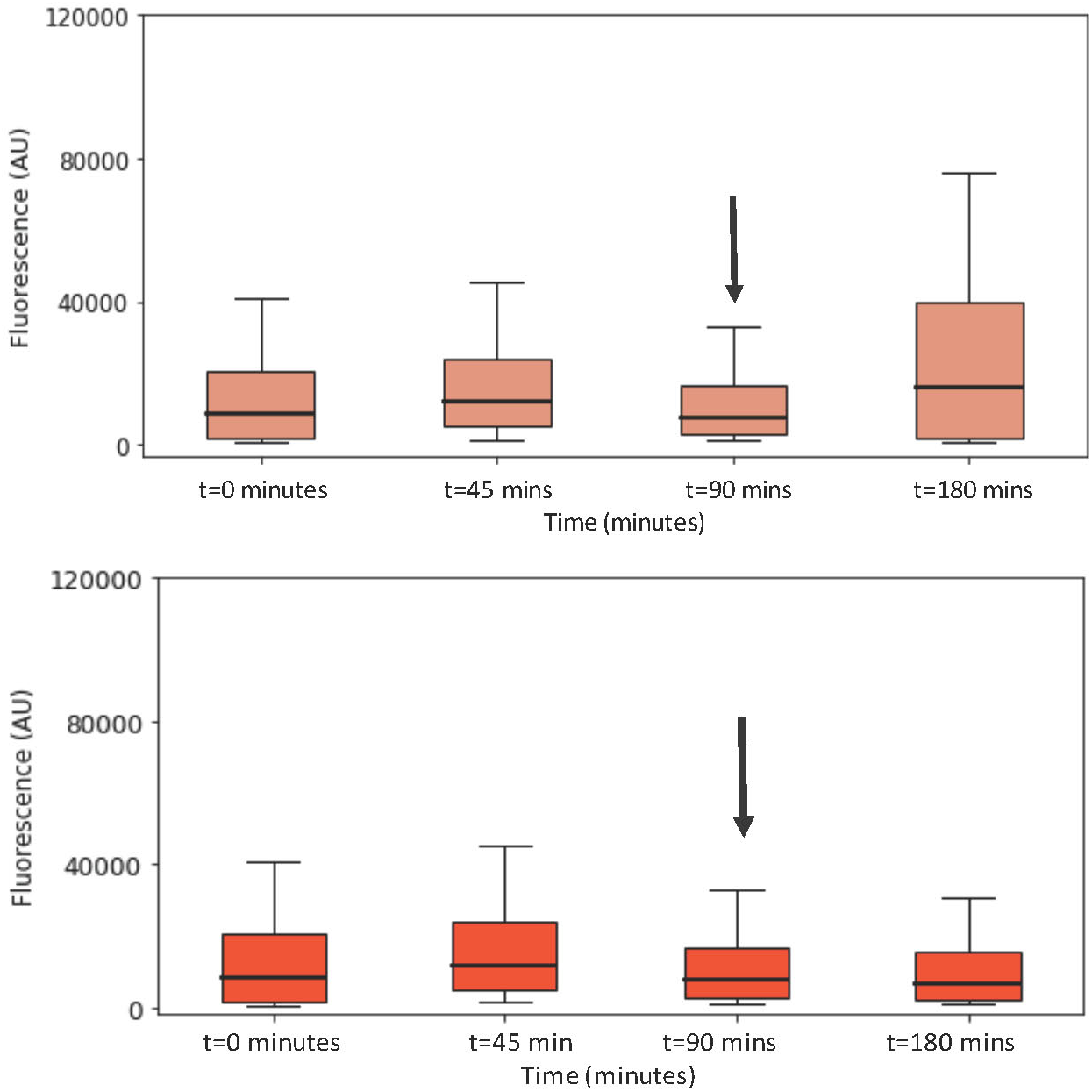
MGA fluorescence can be enhanced by blocking E. coli transcription and promoting T7 induced transcription. MGA6-pet28a circuits were induced by adding 0.1 mM IPTG. They were maintained in their log phase of cell growth (with serial dilution) during 90 mins. Then, they were diluted (arrow in the figure) into fresh M9CA media containing 0.1 mM IPTG (red), and 0.1 mM IPTG and 0.5 mg/ml rifampicin (light red).

The fluorescence signals of the MGA scaffolds can be dynamically controlled by turning off the transcription of MGA. Removal of IPTG from the growth media by washing and suspending the cells in growth media without IPTG was seen to sharply decrease MGA fluorescence to the basal level within 1.5 hours (Figure 3), a time interval that corresponds to approximately two growth cycles (Figure S2). This suggests that dilution due to cell doubling causes the reduction of the signal and more importantly that the MGA-MG signal half-lifetime is at least larger than the cell doubling time similar to that of Spinach(17).

**Figure 3:**
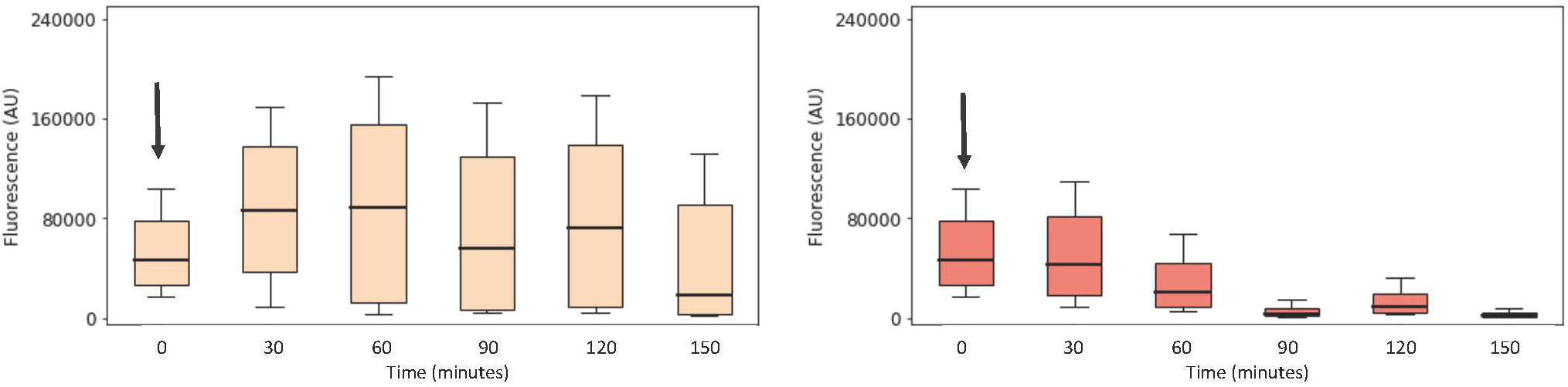
MGA fluorescence can be eliminated by removing IPTG from growth media. MGA fluorescence (MGA6-pET28a) was induced by adding 0.1 mM IPTG M9CA and maintained the log phase of cell growth (with serial dilution) by diluting every 45 minutes. The fluorescence-induced cells were subsequently washed at t=0 (arrow) and diluted into fresh M9CA media in the presence of IPTG (0.1 mM IPTG, right) or its absence (left).

### Robust MGA fluorescence can be seen in the presence of fluorescent proteins

One advantage of MGA is its fluorescence spectrum at far-red wavelength that allows it to be used in conjugation with fluorescent proteins and aptamers in green and red wavelengths. To study the possibility of observing fluorescent protein and MGA fluorescence simultaneously, MGA6 was inserted in a high copy number plasmid with pMB1 origin (pGA1A3) containing a single gene expression cassette driven by *lac*-promoter, which regulates the expression of both MGA6 and a gene encoding for enhanced yellow fluorescent protein (eYFP; BBa_E0030; stop codon included; Figures 4 and 5) or enhanced green fluorescent protein (eGFP; BBa_E0040; stop codon included; Figure S3). Upon induction by IPTG, a fusion RNA consisting of MGA6 and the fluorescent protein mRNA is transcribed. Eventually only the mRNA component of the fusion RNA is further translated into the fluorescent protein. The fluorescence signals shown in Figure 4(B-D) correspond to the fusion mRNA(eYFP)-MGA6 and eYFP.

**Figure 4:**
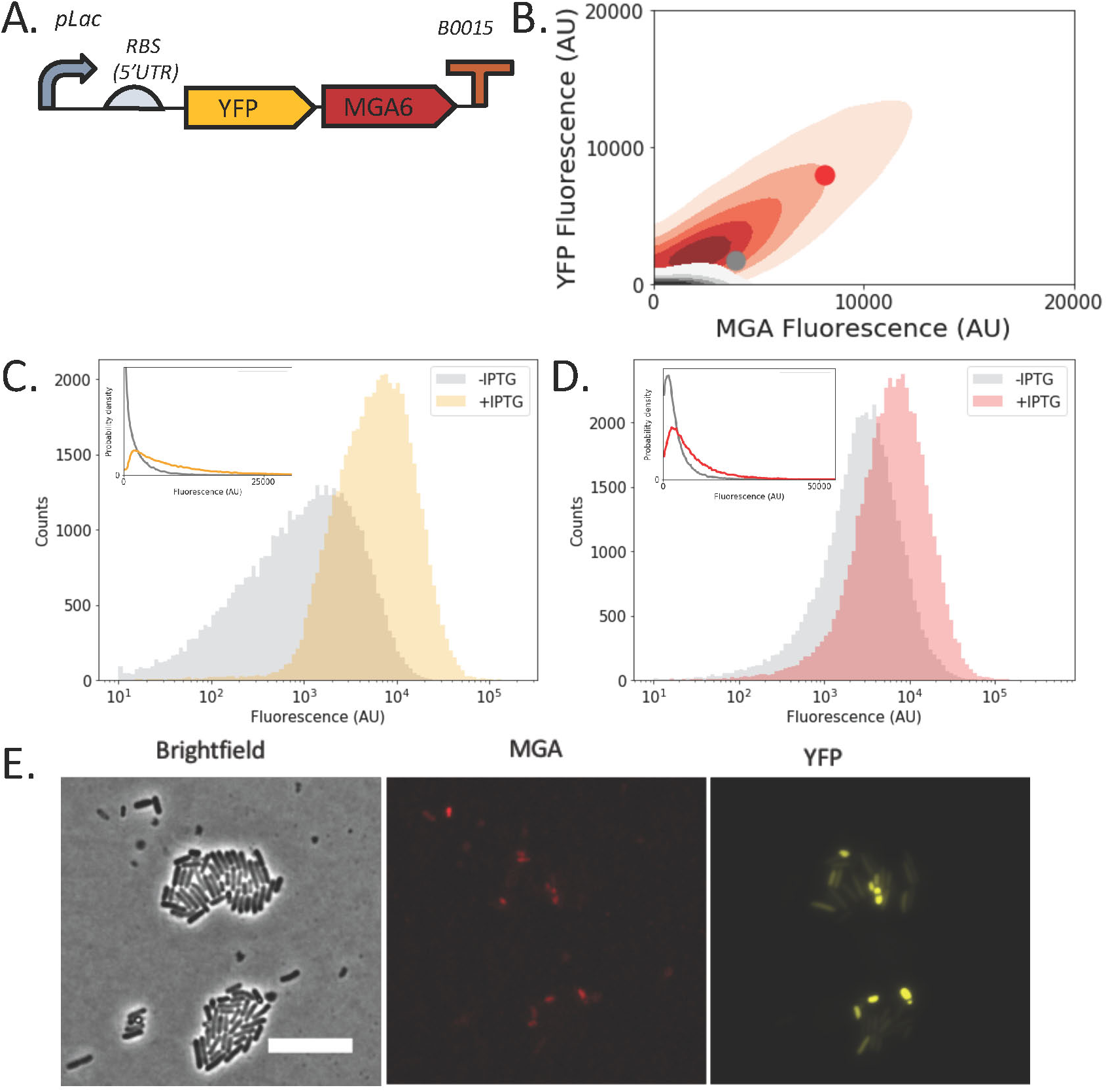
Inducible co-expression of MGA and eYFP. (A) The MGA6-eYFP fusion expression is regulated by plac that can be induced with IPTG. The gene expression cassette is placed in pGA1A3. RBS denotes ribosome binding site. (B-E) MGA-MG fluorescence was measured in response to 5 µM MG along with 0.1 mM IPTG, for cells growing in M9CA media. MGA-MG and eYFP fluorescence were quantified using a flow cytometer (B, C, D) and a microscope (E). (B) Two dimensional density plot of MGA and eYFP signals show positive correlation between the two for both the uninduced (-IPTG, gray) and the induced (+IPTG, red). SBOL compliant figure (31) of the construct is shown (A). Scale bar in (E) corresponds to 10 µm.

**Figure 5:**
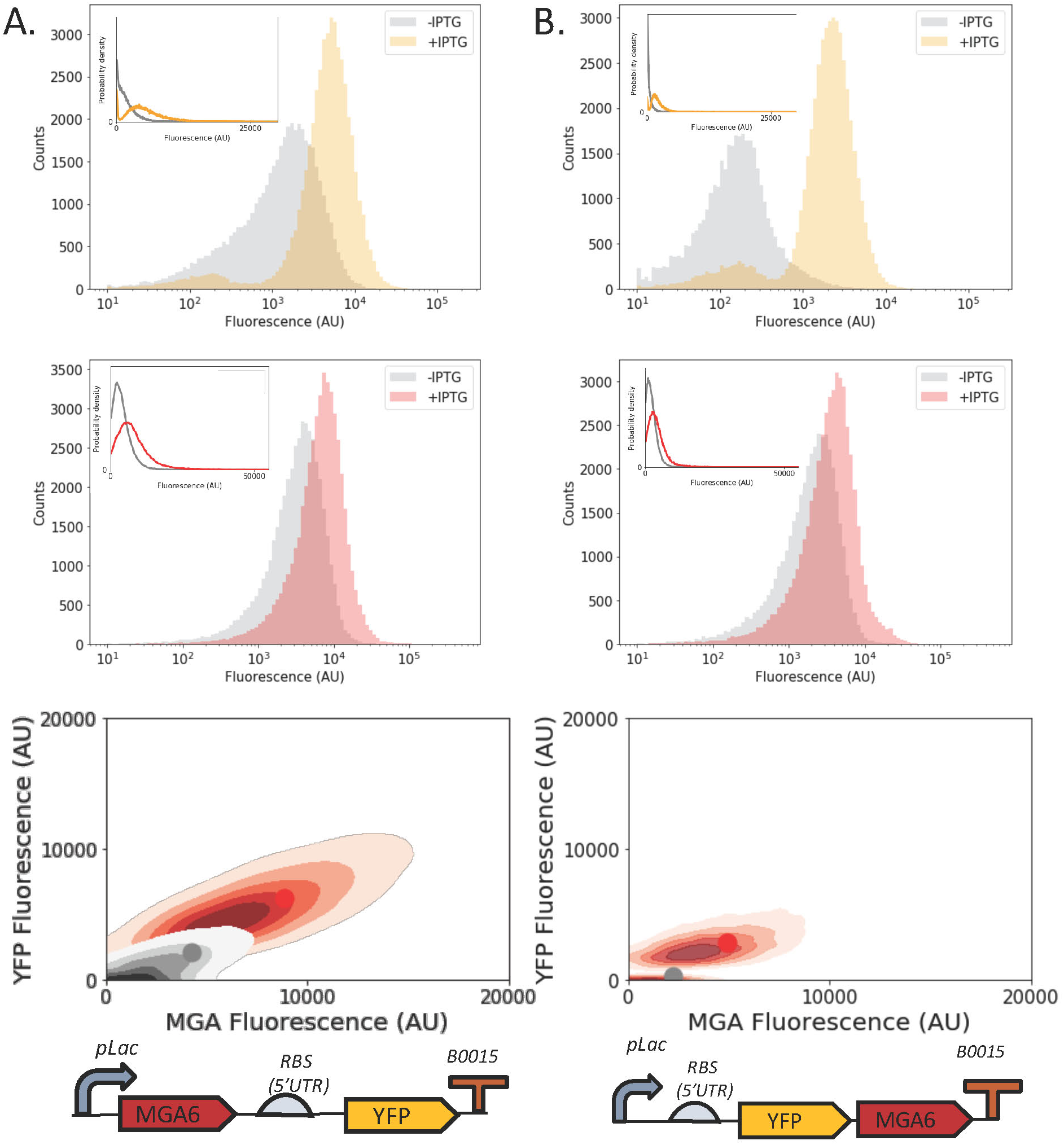
Inducible co-expression of MGA when placed at 5’ or 3’ UTR of eYFP. This figure shows MGA and eYFP fluorescence in induced and un-induced *E. coli* cells containing a fusion eYFP-MGA6 and MG6-eYFP in response to 5 µM MG. MGA expression was induced by adding 0.1 mM IPTG to cells growing in M9 media and is quantified using a flow cytometer. The MGA6 and eYFP fluorescence signal distributions from individual cells and SBOL compliant figures of the construct are shown.

At the single cell level, we confirmed that a strong correlation between the levels of MGA and eYFP via flow cytometry (Figure 4B) and microscopy (Figure 4E). These results indicate the potential of using MGA as genetically encoded fluorescent tags for RNA molecules including in principle both mRNAs and regulatory RNAs to study their roles.

### Robust MGA fluorescence can be seen in the presence of other aptamers

Recent *in-vivo* studies in *E. coli* involving the expression of aptamers such as Spinach and Mango have shown promising results(17,18,23-27). The fact that fluorescence wavelengths from these aptamers are respectively green and yellow show that they could be used simultaneously with MGA’s far-red fluorescence (630/655 nm) for the simultaneous labeling or color-coding of two different RNAs. A fusion of MGA scaffolds and Spinach (469/501 nm)(23) or Mango (510/535nm)(25) scaffolds, separated by linkers, were constructed in a pET28a plasmid. *E. coli* expressing these cells exhibited a large increase in MGA fluorescence but a moderate increase in Spinach (64 repeats) or Mango (2 repeats) fluorescence upon induction (Figure S4). The moderate increase in Mango or Spinach can be perhaps due to the interference from the MGA scaffolds, or inserted linker sequences used for Spinach or Mango scaffolds. While further experimental optimization for Spinach or Mango is required, this experiment shows that the MGA scaffolds (4 repeats) are robust when fused with other types of RNA aptamer probes.

In summary, expression of MGA scaffolds in *E. coli* exhibited a strong, stable MGA fluorescence that could be quantified via flow cytometry and microscopy. The far-red fluorescence of MGA can be used in conjunction with a large variety of fluorescent proteins or aptamers in green or yellow wavelengths.

## Discussion

MGA is one of the first light-up aptamers to be described in the literature(12). However comparatively few studies have been conducted on MGA due to a variety of concerns that include high background to fluorescence ratio and cytotoxicity. Recent studies on other light-up aptamers such as Spinach(17), RNA Mango(25), Spinach 2(24), Broccoli(26), *etc.* have revealed various methodologies and techniques that can also be applied to the MGA system to overcome its shortcomings that will enable its use as a RNA labeling and imaging marker. Some of its potential advantages over widely applied RNA labeling methods such as using fluorescent protein-fused RNA binding proteins include low fluorescence background, elimination of separate introduction of RNA binding proteins, and evasion of perturbation on target RNAs by protein binding. Additionally, the far-red fluorescence of MGA can be used in conjugation with other light-up aptamers which emit light in green and yellow wavelengths such as Spinach, Spinach2 and Mango(23-25). Another advantage of MGA is the low cost and availability of MG compared to Spinach and Mango.

The recent study on Spinach aptamer(17) demonstrated that a scaffold of multiple Spinach aptamer repeats displayed a more robust aptamer-induced fluorescence that allowed better RNA visualization in bacterial cells. Here, we applied these findings to design a scaffold of multiple MGA repeats and express them in bacterial cells using an inducible plasmid. We observe a strong and robust MGA fluorescence, showing more than a 20-fold increase in MGA fluorescence upon induction. These results present a significant improvement over previous published results that show only up to 3-fold increase in fluorescence in batch cultures(15).

Previous studies have shown that MG concentrations below 10 µM do not have any effect on *E. coli* viability, even though MG toxicity has been one of the concerns of using MGA *in vivo(28)*. For this paper, we have used malachite green concentration of 5µM to study MGA induced fluorescence. MGA fluorescence can be quantified by adding much lower levels of MG (5µM) than the amounts of DFHBI (100 µM) used in spinach fluorescence in previous studies in *E. coli*(17,23). This can be partly because the K_d_ of MGA to MG dye (K_d_=0.11 µM) is lower than that of Spinach to DFHBI (K_d_=0.3 µM).

Previous studies on both Spinach(29) and MGA(15) have shown that placing an aptamer in either the 5’ or 3’ untranslated region (UTR) of a gene of interest decreases the fluorescent output of the aptamer possibly due to increased misfolding rates through interactions with proximal nucleotides. However, using our MGA scaffolds we show that the fluorescence is not diminished in the presence of eYFP, eGFP, Spinach, or Mango.

A recent study on a similar scaffold of multiple MGA repeats in eukaryotic cells indicated that the MGA system can only be used for about 10 minutes after the addition of MG dye where the fluorescence/background levels remain sufficiently low(18). However, using MGA6 we observe a decreased yet detectable (approximately two-fold above the basal level) fluorescence even in 6 hours after the addition of MG to the growth media (Figure S5). MGA fluorescence relies on the specific folding of MGA upon its binding to MG. It is to be noted that our MGA6 has different designs and different linkers (designed to maintain MGA folding) when compared with the MGA repeats used in the other study(18) which might impact RNA folding processes differently.

One main advantage of MGA6 is that it is smaller (429 bp) than fluorescent proteins or 8 repeats of Spinach aptamer. The smaller MGA4 (297 bp) or MGA-5S (189 bp) have also shown to be capable of a moderately strong fluorescence. A recent paper identified new analogs of MGA that are capable of binding to MG and induce fluorescence. One such analog, MGA 1.3 (26 bp) is smaller in size than the 38 bp core MGA that we used here. This analog can be potentially used to design MGA scaffolds to decrease size (30).

In conclusion, our scaffold of multiple MGA repeats was shown to be capable of generating strong MGA-MG fluorescence in *E. coli* under various growth conditions. This strong MGA fluorescence was not diminished when fused with fluorescent protein mRNAs or RNA aptamers. This study demonstrates an enhanced fluorescence quality of the MGA scaffold that can potentially be used for live cell studies using flow cytometry and microscopy. In addition, it will enable dual color monitoring of multiple RNA molecules, which can be used for direct understanding of RNA regulation and transcription-translation.

## Methods

### Strains and media

Turbo Competent *E. coli* (NEB) and LB media (Sigma) were used for plasmid preparation. All studies were performed using the *E. coli* strain BL21DE3 (NEB) or 10B (NEB). Cells used in experimental studies were cultured in M9 Minimal media (Teknova, 42mM Na_2_HPO_4_, 24mM KH_2_PO_4_, 9mM NaCl, 19mM NH_4_Cl), 1mM MgSO_4_, 0.1mM CaCl_2_, 2.0% glucose, 0.5µg/ml thiamine) or with M9CA media (Teknova, 42mM Na2HPO4, 24mM KH_2_PO_4_, 9mM NaCl, 19mM NH4Cl), 0.2mM MgSO_4_, 0.1mM CaCl2, 1.0% glucose, 0.5µg/ml thiamine, 0.1% casamino acids). All media was supplemented with antibiotics, 100 µg/ml Carbenicllin (Teknova) and 50 µg/ml Kanamycin (Teknova), and inducer, Isopropyl β-D-1- thiogalactopyranoside (IPTG; Sigma), when appropriate.

### Plasmid construction

A scaffold of 6 MGA repeats (MGA6) was designed using the MGA sequence described in the literature(15). The MGA repeats were separated by randomized 17 nt linker sequences and two complementary base pair sequences at both ends of MGA that stabilize the MGA structure. Predicted RNA folding determined by RNAfold(19) and Kinefold(20) was used in the design process to select for linkers that do not disrupt MGA native folding.

The MGA6, MGA12 and MGA-5S sequences were synthesized as double-stranded DNA and was delivered in the pUC57 plasmid (Genscript) between the XbaI and HindIII restriction sites (pUC57-MGA6) by Genscript. The scaffolds were excised with XbaI and HindIII and inserted into pET28a plasmid between the XbaI and HindIII restriction sites to generate pET28a-MGA6.

The MGA6 sequence was amplified pUC57-MGA6 using inserted PCR amplification and Phusion DNA polymerase (NEB). A pGA1A3 plasmid containing a synthetic biology circuit consisting of a plac promoter, a ribosome binding site (RBS), fluorescent protein and a terminator was similarly linearized using PCR amplification using Phusion DNA polymerase with primers that share overlap sequences with the MGA6 amplicon. The MGA6 and a backbone amplicons were ligated in the Gibson assembly method by using a one-step isothermal DNA assembly mixture that includes Phusion DNA Polymerase, T4 Ligase (NEB), and T5 Exonucleases (NEB). pGA1A3 is a high copy number plasmid with ampicillin resistance and a pMB1 origin of replication. The resulting plasmid DNA was transformed into chemically competent Turbo cells (NEB) and plated on selective LB agar.

### Flow cytometry

Cells were inoculated into M9 media and grown overnight at 37°C and 250 r.p.m. shaking. From this seed culture, the cells were then diluted to a 1:10000 ratio and incubated at 37°C grown to OD600=0.05-0.1 in a shaker. MGA fluorescence was measured in a Sony Biotechnology ec800 flow cytometer using a 665 nm filter (equipped with a 660 nm splitter) and a 640 nm excitation laser. 50,000 events were collected for each sample per run and gated by using a 2-D normal distribution using python package FlowCytometryTools. A Medium Flush cycle was used after each run to avoid inter-sample contamination.

### Fluorescence microscopy

Imaging was performed using a Nikon Ti-E inverted wide-field fluorescence microscope with a large-format scientific complementary metal-oxide semiconductor camera (NEO, Andor Technology) and controlled by NIS-Elements. Samples were kept at 30°C throughout the imaging process using an environmental chamber. The total imaging time for each experiment was 10 hours during which both brightfield and fluorescent images were captured every 3 minutes. Image processing and analysis was completed using ImageJ (NIH)(21).

Prior to measurement, cells were suspended in warm M9 media supplemented with 5 µM malachite green. 5 µl of suspended culture was placed on 0.5% M9 agar containing 5 µM malachite green. A glass microscope slide was placed over the agar pad and sealed airtight with a 1:1:1 mixture of Vaseline, lanolin, and paraffin.

## Supporting Information

Supporting Note: Five supplementary figures and DNA sequences of our MGA scaffolds.

## Author Information

Kyung Hyuk Kim: Department of Bioengineering, University of Washington, Seattle, Washington 98105, United States

V. Siddartha Yerramilli: Department of Bioengineering, University of Washington, Seattle, Washington 98105, United States

## Author Contribution

KHK designed and managed the project, performed preliminary experiments, and wrote the manuscript. VSY executed the project, designed MGA scaffolds, performed their experimental characterization, and wrote the manuscript.

## Acknowledgements

We would like to thank Herbert Sauro for providing us with the lab space and reading the manuscript, Peter Unrau for providing us the Mango aptamer sequence and TO-1 dye, and Taekjip Ha for providing us with Spinach aptamer sequence. We would also like to thank Joo-Young Lee, James Carothers, and Paul Wiggins for their advice and suggestions. We would also thank Dr. Wiggins and his lab for allowing us to use the microscope. This work has been supported by National Science Foundation (Grant number MCB 1515280 to K.K).

## Supplementary Materials

**Figure S1:**
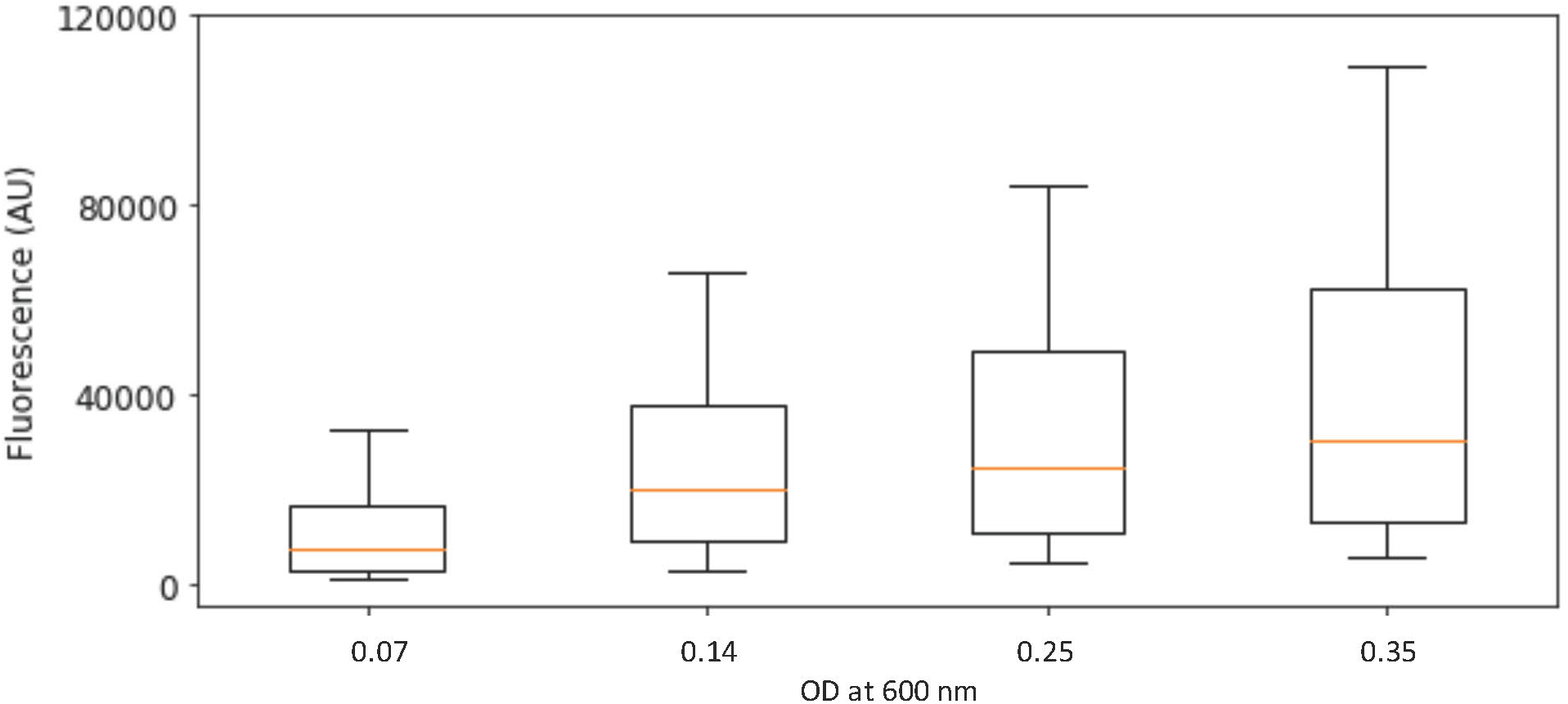
Inducible malachite green aptamer expression: This figure shows MGA fluorescence in induced (+IPTG) *E. coli* BL21DE3 cells containing a pET28a plasmid consisting of a MGA6 scaffold in response to 5µM malachite green at various values of OD600. MGA expression was induced by adding 0.1 mM IPTG to cells growing in M9CA media and is quantified using a flow cytometer.

